# Plexin-B1 and Plexin-B2 play non-redundant roles in GABAergic synapse formation

**DOI:** 10.1101/2023.10.27.564428

**Authors:** Susannah S. Adel, Zachary J. Pranske, Tess F. Kowalski, Nicole Kanzler, Roshni Ray, Catherine Carmona, Suzanne Paradis

## Abstract

Synapse formation in the mammalian brain is a complex and dynamic process requiring coordinated function of dozens of molecular families such as cell adhesion molecules (CAMs) and ligand-receptor pairs (Ephs/Ephrins, Neuroligins/Neurexins, Semaphorins/Plexins). Due to the large number of molecular players and possible functional redundancies within gene families, it is challenging to determine the precise synaptogenic roles of individual molecules, which is key to understanding the consequences of mutations in these genes for brain function. Furthermore, few molecules are known to exclusively regulate either GABAergic or glutamatergic synapses, and cell and molecular mechanisms underlying GABAergic synapse formation in particular are not thoroughly understood. However, we previously demonstrated that Semaphorin-4D (Sema4D) regulates GABAergic synapse development in the mammalian hippocampus while having no effect on glutamatergic synapse development, and this effect occurs through binding to its high affinity receptor, Plexin-B1. Furthermore, Plexin-B2 contributes to GABAergic synapse formation as well but is not required for GABAergic synapse formation induced by binding to Sema4D. Here, we perform a structure-function study of the Plexin-B1 and Plexin-B2 receptors to identify the protein domains in each receptor that are required for its synaptogenic function. We also provide evidence that Plexin-B2 expression in presynaptic parvalbumin-positive interneurons is required for formation of GABAergic synapses onto excitatory pyramidal neurons in CA1. Our data reveal that Plexin-B1 and Plexin-B2 function non-redundantly to regulate GABAergic synapse formation and suggest that the transmembrane domain may underlie these functional distinctions. These findings lay the groundwork for future investigations into the precise signaling pathways required for synapse formation downstream of Plexin-B receptor signaling.

## Introduction

Proper circuit function in the mammalian brain relies upon cell-cell communication through two main types of chemical synapses: glutamatergic (excitatory) and GABAergic (inhibitory). In the cortex, activity of excitatory pyramidal neurons is tightly controlled by inhibitory interneurons, the dysfunction of which has been linked to disorders such as autism and epilepsy (Ferguson & Gao, 2018; Takano & Sawai, 2019). Therefore, it is critical to understand mechanisms by which GABAergic interneurons form the proper synaptic connections with both glutamatergic pyramidal neurons and GABAergic interneurons.

Synapse formation is a complex process occurring in several steps. First, initial contact between a pre- and postsynaptic neuron occurs, followed by adhesion and stabilization of this point of contact. Subsequently, synaptic machinery accumulates in both the presynaptic active zone and postsynaptic specialization, specifically: neurotransmitter release machinery, scaffolding proteins, neurotransmitter receptors, and other required signaling molecules (reviewed in Batool et al., 2019; McAllister, 2007; Scheiffele, 2003). Numerous families of secreted or membrane attached ligand/receptor pairs and cell adhesion molecules (CAMs), expressed in neurons and in glia are implicated in synapse development and function (Sudhof, 2018). While it is well-established that molecules such as Ephs/ephrins, neuroligins/neurexins, Nrg1/ErbB4, FGF/FGFR, CAMs, and Semaphorins/Plexins regulate synapse formation, it is an outstanding challenge to determine precisely which steps of synapse formation these molecules control, as well as whether they exert their functions from the presynaptic or postsynaptic neuron or both (Craig & Kang, 2007; Dabrowski et al., 2015; Hruska & Dalva, 2012; Koropouli & Kolodkin, 2014; Luo et al., 2021; Sudhof, 2018; Terauchi et al., 2010).

Importantly, it has also been a challenge to define mechanisms that differ between glutamatergic and GABAergic synapse formation, in part because few synaptogenic molecules have been identified that selectively promote formation of GABAergic, but not glutamatergic, synapses. One exception to this is that Sema4D signaling through its receptor Plexin-B1 regulates GABAergic, but not glutamatergic, synapse formation. We previously demonstrated that application of the extracellular domain of Sema4D either in vitro or in vivo caused the rapid formation (2hrs) of functional, GABAergic synapses (Adel et al., 2023; Kuzirian et al., 2013). Using a live imaging approach, we showed that Sema4D application affects the behavior of GFP-labeled gephyrin which is a postsynaptic scaffolding protein at GABAergic synapses. Taken together, our data favor a model whereby Sema4D/Plexin-B1 signaling acts at the recruitment step in synapse formation to localize synaptic components to the nascent synapse.

In addition, recent advances from our group demonstrated the therapeutic potential of targeting GABAergic synapse formation and function in the case of neurological disorders involving hyperexcitability, underscoring the importance of delineating these processes (Acker et al., 2018; Adel et al., 2023). For example, we previously found that application of the Sema4D extracellular domain to neuronal cultures increased GABAergic synapse density within 30 minutes, increased inhibitory tone within 2hrs of application, and increased resistance to seizure in various rodent models of epilepsy (Adel et al., 2023; Kuzirian et al., 2013). The selective, rapid, and functional effects of Sema4D motivated our further investigation into the mechanisms of synapse development mediated by Semaphorin/Plexin signaling.

Semaphorins, a 25-member family of secreted and membrane-attached ligands containing a conserved, extracellular Sema domain, bind to the extracellular Sema domain of Plexin receptors (reviewed in Alto & Terman, 2017; Koropouli & Kolodkin, 2014; Tamagnone & Comoglio, 2000; Yazdani & Terman, 2006). Plexins are a family of 9 transmembrane receptors with a wide range of functions including regulation of cell migration and proliferation (reviewed in Tamagnone et al., 1999; Toledano & Neufeld, 2023). Class 4 Semaphorin dimers bind to Plexin-B receptors to regulate numerous functions in the developing nervous system, including neural tube closure, cerebellar cell migration and differentiation, and actin cytoskeleton remodeling (Deng et al., 2007; Oinuma et al., 2010; Oinuma et al., 2003; Worzfeld et al., 2014).

In the past decade, we established novel roles for other Class 4 Semaphorins and their Plexin-B receptors in synapse formation. Both Sema4A and Sema4D promote GABAergic synapse formation via signaling through the Plexin-B1 receptor (Kuzirian et al., 2013; McDermott et al., 2018). Additionally, we showed that another Plexin-B family member, Plexin-B2, is required in the postsynaptic neuron for proper GABAergic synapse formation (McDermott et al., 2018). Interestingly, while there is no evidence to suggest that Plexin-B1 mediates glutamatergic synapse formation, signaling by Sema4A with Plexin-B2 drives an increase in the density of glutamatergic synapses in dissociated hippocampal neurons. These findings suggest a model whereby Plexin-B1 and Plexin-B2 exhibit distinct synaptogenic roles. In situ hybridization revealed that Plexin-B1 and Plexin-B2 are expressed in both presynaptic inhibitory interneurons and in postsynaptic excitatory neurons in the hippocampus (McDermott et al., 2018), suggesting that differential expression is unlikely to explain functional distinctions between these molecules.

Here, we sought to reveal distinct functions of Plexin-B1 and Plexin-B2 receptors in GABAergic synapse formation and uncover their underlying molecular mechanisms. First, we interrogate the role of Plexin-B2 expressed in parvalbumin-positive (PV) cells in the hippocampus in GABAergic synapse formation. We next ask if Plexin-B1 and Plexin-B2 function redundantly in GABAergic synapse formation. Finally, we perform a structure-function analysis to identify roles for distinct Plexin-B1 and Plexin-B2 molecular domains in driving formation of GABAergic synapses. Our data reveal unique roles for different Plexin-B receptor domains in GABAergic synapse formation and provide new insights into the precise signaling configurations by which Plexin-B receptors mediate this process.

## Results

### Plexin-B2 is expressed in PV+ interneurons

In the hippocampus, Plexin-B1 and Plexin-B2 are expressed in interneurons and in excitatory pyramidal neurons (McDermott et al., 2018). We previously provided evidence that Sema4D expressed in non-neuronal cells promotes clustering of presynaptic molecular components of GABAergic synapses when co-cultured with neurons. Importantly, this effect required Plexin-B1 expression in the presynaptic neurons (McDermott et al., 2018). Here, we sought to determine if Plexin-B2 expression in the presynaptic neuron is also required for GABAergic synapse formation. The identity of interneuron subclasses that express Plexin-B2 is unknown. Thus, we first assessed whether Plexin-B2 is expressed in PV+ interneurons. We isolated and fixed brains from wild-type mice at P14 and performed single molecule in situ hybridization to probe for parvalbumin (*Pvalb*) mRNA and Plexin-B2 (*Plxnb2*) mRNA in the hippocampus. We validated the in situ hybridization protocol using both positive and negative control probes (Fig. S1). We examined select hippocampal subregions and quantified the percentage of *Pvalb*-expressing cells that also expressed *Plxnb2*. We found that *Plxnb2* mRNA is expressed in greater than 70% of *Pvalb-*positive cells across all subregions (Fig. 1A,B).

**Figure 1.**
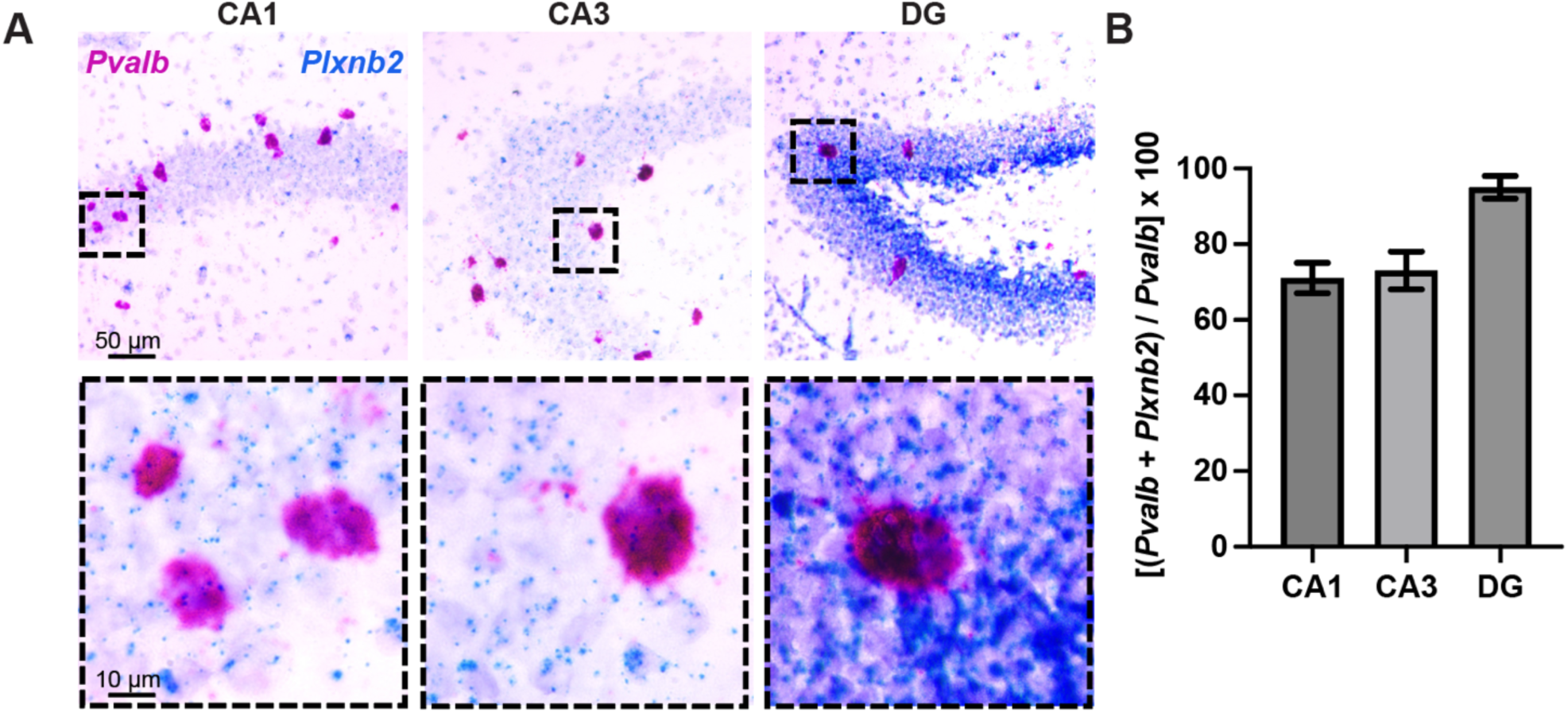
*Plxnb2* mRNA is expressed in *Pvalb*-positive interneurons in the hippocampus. (A) Representative images of mRNA expression in the indicated hippocampal subregions in tissue isolated from P14 mice (*Plxnb2* in blue, *Pvalb* in red, hematoxylin counterstain in light purple). Insets indicated by dashed boxes are shown at higher magnification in bottom panels. (B) Quantification of the percentage of *Pvalb*+ cells expressing *Plxnb2* mRNA (*Plxnb2* + *Pvalb*) divided by the total number of *Pvalb*+ cells in the indicated subregions. n = 2 sections from two animals. Error bars indicate SEM for binary distribution.

### Plexin-B2 is required in PV+ interneurons for proper GABAergic synapse development

PV+ neurons critically modulate the activity of excitatory pyramidal neurons in the hippocampus via inhibitory synaptic connections onto the cell soma perisomatic area of pyramidal neurons (Ferguson & Gao, 2018). Thus, we next sought to determine whether Plexin-B2 expression in PV+ interneurons mediates the formation of GABAergic synapses onto these excitatory pyramidal neurons. We crossed mice harboring a transgene driving expression of Cre recombinase in PV-positive cells (*PV^Cre^)* with Plexin-B2 cKO (*Plxnb2^flx/flx^*) mice to obtain a *PV^Cre^;Plxnb2^wt/flx^*mouse line. We confirmed the expression of Cre in PV+ interneurons in this line by performing an intracerebroventricular injection of Cre-dependent AAV.FLEX.TdTomato virus into pups aged P1-P3. At approximately 3 months of age, we processed the mice for immunohistochemistry using antibodies recognizing PV and GAD65, a protein localized exclusively to GABAergic presynaptic boutons, and acquired images on a confocal microscope (Fig. 2A).

**Figure 2.**
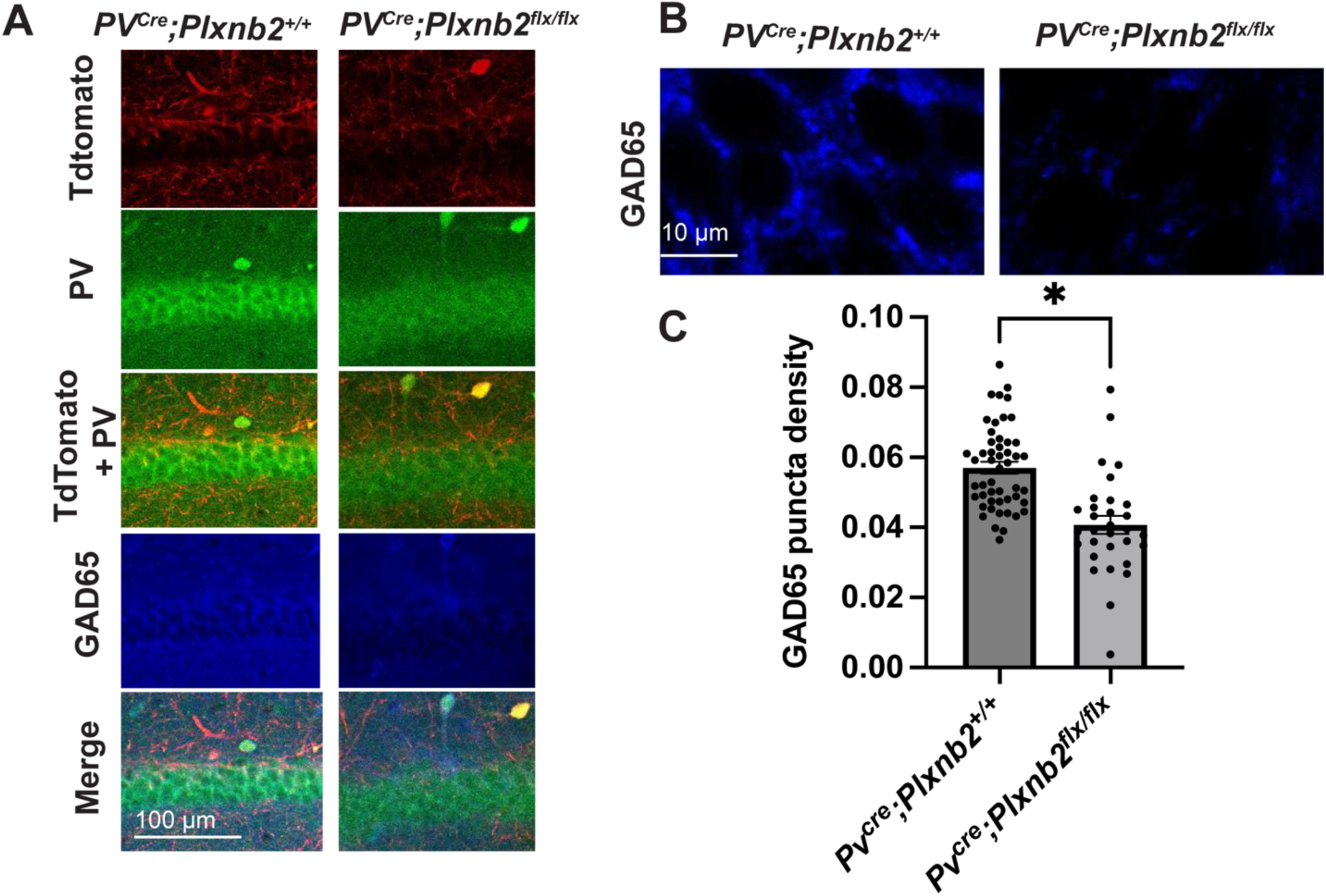
Plexin-B2 expression in presynaptic PV+ interneurons is critical for GABAergic synapse formation. (A) Representative images of CA1, tissue sections indicate expression of the Cre-dependent TdTomato cell fill (red) delivered by ICV injection into *PV^Cre^*;*Plxnb2^flx/flx^*mice at P1-3, as well as immunostaining for PV (green) and GAD65 (blue). (B) Representative ROIs in CA1 in which GAD65 (blue) puncta density was quantified. (C) Quantification of number of GAD65+ puncta/ area of soma, normalized to the area of the ROI. *p < 0.05, unpaired t-test. N = 31-50 ROIs from 3-8 mice per condition. Error bars are ± standard error of the average ratio. Circles represent individual ROIs.

PV+ interneurons predominantly form synaptic contacts onto the perisomatic region of excitatory pyramidal neurons (Ferguson & Gao, 2018). Therefore, we drew regions of interest of approximately equal size in the stratum pyramidale layer in the CA1 region of hippocampus where the somas of these pyramidal neurons are located (excluding areas that were positive for PV signal) and quantified the density of GAD65 puncta in each ROI (GAD65 immunoreactivity is a reliable proxy for GABAergic synapse density). This analysis revealed decreased GAD65 density in the hippocampus of *PV^Cre^;Plxnb2^flx/flx^*mice as compared to controls (*PV^Cre^;Plxnb2^+/+^*) (Fig. 2A-C). Based on this result we conclude that Plexin-B2 expressed in presynaptic PV+ interneurons is required for normal GABAergic synapse formation onto pyramidal neurons in the hippocampus.

### GABAergic synapse formation onto PV+ interneurons is unaffected by Plexin-B2 knockout

Given the role of Plexin-B2 in pyramidal neurons to mediate GABAergic synapse formed onto these neurons, we sought to determine if Plexin-B2 also mediated synapse formation onto interneurons. PV+ cells receive contacts from both excitatory and inhibitory neurons, including from SST+ and VIP+ interneurons and a smaller number of inputs from PV+ neurons (reviewed in Cardin, 2018). We used the same images analyzed in Figure 2 to determine if Plexin-B2 expressed in PV+ cells regulates GABAergic synapse formation onto PV+ cells themselves. We drew ROIs around cell somas that were both PV+ and expressing Cre-dependent TdTomato, indicating they were both PV+ and had Plexin-B2 deletion, in or near the pyramidal cell layer of hippocampus and then quantified the density of GAD65 puncta onto the soma. All cell bodies within CA1, CA2, or CA3 that fit these criteria were analyzed. We found no significant difference in the GAD65 puncta density onto PV+ interneurons in tissue from *PV^Cre^;Plxnb2^+/+^* and *PV^Cre^;Plxnb2^flx/flx^* mice, suggesting that Plexin-B2 expression in PV+ interneurons does not regulate the development of perisomatic GABAergic synapses formed onto these cells (Fig. 3A,B).

**Figure 3.**
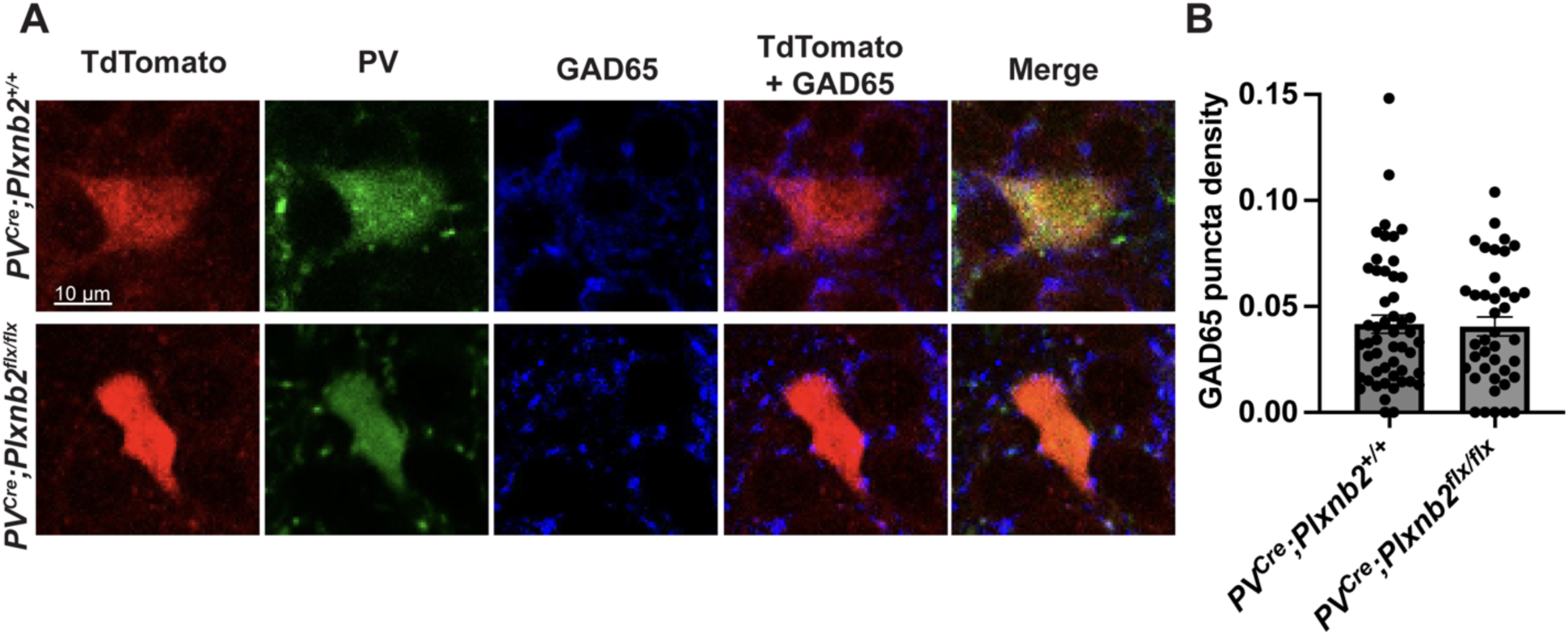
Plexin-B2 is not required for formation of GABAergic inputs onto PV+ interneurons. (A) Representative images of single PV+ neuron somas in CA1 from 3 month old *PV^Cre^;Plxnb2^+/+^*or *PV^Cre^;Plxnb2^flx/flx^* mice that were given an ICV injection of AAV.FLEX.TdTomato between age P1-P3, immunostained with antibodies detecting PV (green) and GAD65 (blue). (B) Quantification of the number of GAD65 puncta divided by the area of the PV+ interneuron cell soma. No significant differences were detected by an unpaired t-test. Circles represent individual cells.

### Plexin-B1 and Plexin-B2 do not function redundantly in GABAergic synapse formation

Previously we showed that RNAi-mediated Plexin-B2 knockdown in cultured hippocampal neurons caused a decrease in GABAergic synapse density while Plexin-B1 knockdown did not (McDermott et al., 2018). This result was puzzling when considered in combination with our data derived from experiments using a *Plxnb1^−/−^* mouse strain, demonstrating a requirement for Plexin-B1 to mediate Sema4D-dependent GABAergic synapse formation (Kuzirian et al., 2013). We hypothesized that compensation from Plexin-B2 in the absence of Plexin-B1 might explain these findings. In support of this hypothesis, a 2016 study showed that double deletion of Plexin-B1 and Plexin-B2 resulted in impaired cortical neurogenesis, despite each single deletion yielding no phenotype (Daviaud et al., 2016).

Thus, we sought to examine the effects on GABAergic synapse formation upon genetic knockout of Plexin-B1 and Plexin-B2 alone or in combination. We reasoned that if Plexin-B2 does in fact compensate for the absence of Plexin-B1, the double knockout condition should exhibit the same or greater deficit in GABAergic synapse density than knockout of either Plexin-B1 or Plexin-B2 alone. To begin, we performed a genetic epistasis experiment using mice of four different genotypes: wild type (WT), Plexin-B1 constitutive knockout, Plexin-B2 conditional KO (cKO), and Plexin-B1/Plexin-B2 double knockout to ask if Plexin-B1 and Plexin-B2 function redundantly to regulate GABAergic synapse development.

To obtain animals with the necessary genotypes, we crossed transgenic mice harboring a Plexin-B1 null allele (*Plxnb1^−/−^*) with mice in which exons 7-9 of the Plexin-B2 gene are flanked by loxP sites (*Plxnb2^flx/flx^*), resulting in excision of exons 7-9 of Plexin-B2 in the presence of Cre recombinase, yielding a null allele (Daviaud et al., 2016). In this manner, we obtained pups with genotypes *Plxnb1*^+/+^;*Plxnb2*^flx/flx^ and *Plxnb1^−/−^*;*Plxnb2*^flx/flx^ (Fig. 4A). When these pups reached age postnatal day 6-10 (P6-P10), we isolated and cultured coronal brain sections containing hippocampus. The following day, we bilaterally infected the hippocampal CA1 region with an AAV serotype 9 virus driving expression of either a Cre-GFP fusion protein or GFP, both downstream of a neuron-specific promoter (human Synapsin I). Thus, the Plexin-B2 gene was knocked out in neurons expressing Cre-GFP recombinase and expressed at endogenous levels in neurons infected with the GFP virus. Importantly, this strategy results in manipulation of Plexin-B2 expression in all neurons where it is endogenously expressed, and genetic manipulation is not restricted to either the pre- or postsynaptic neuron. We cultured the slices for five days after infection before fixing and staining with an antibody recognizing GAD65. Using confocal microscopy, we acquired z-stacks of the stratum pyramidale layer in the CA1 region of the hippocampus. We selected the brightest z-plane for analysis and, after identifying GFP+ cells (indicating successful infection), individual cell somas were identified and traced. Following, we quantified the number of GAD65+ puncta overlapping with each soma.

**Figure 4.**
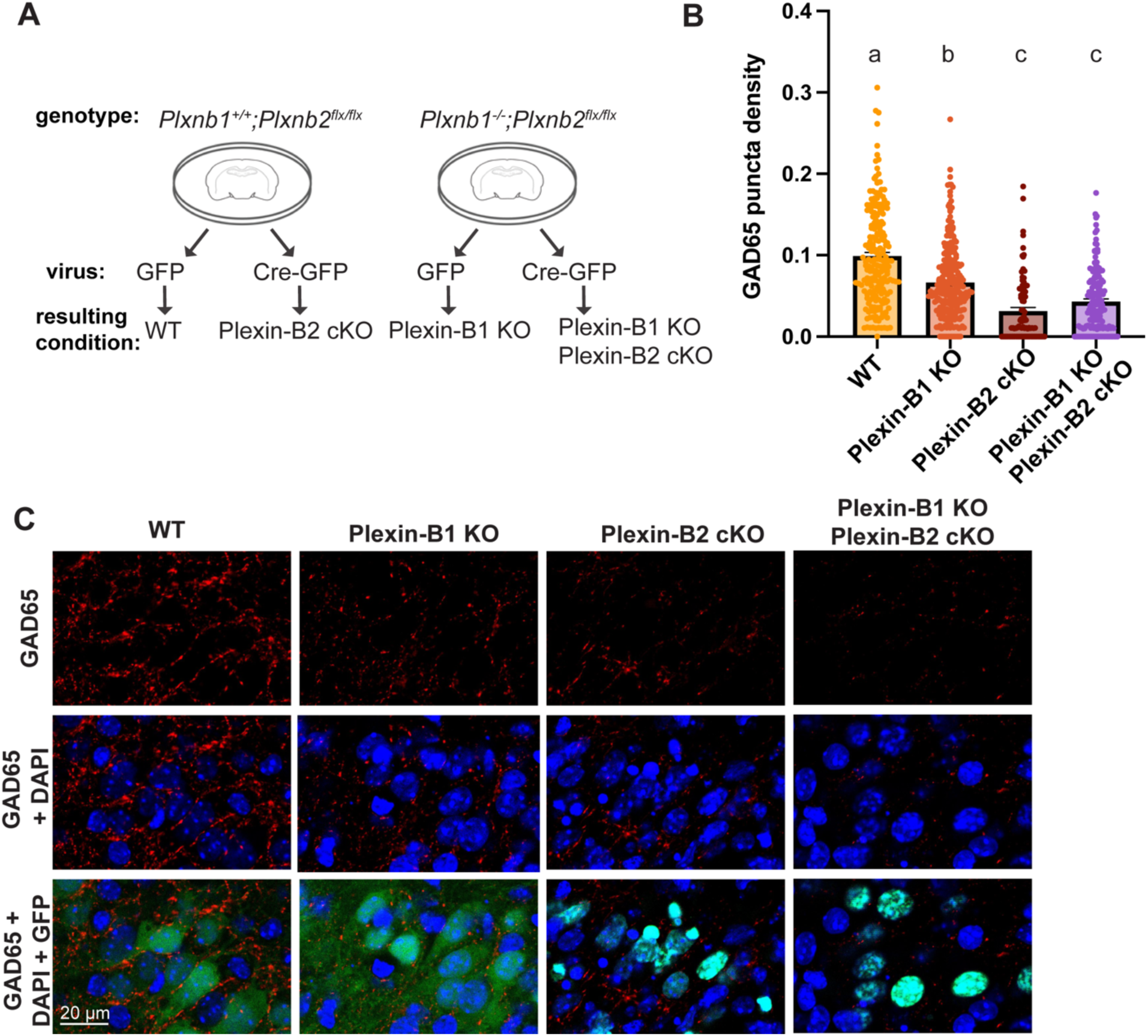
Plexin-B1 and Plexin-B2 do not function redundantly to regulate GABAergic synapse formation. (A) Strategy for gene knockout resulting in single or double deletion of Plexin-B1 and/or Plexin-B2. (B) Quantification of number of GAD65+ puncta divided by area of soma. At least 80 cells were sampled across 5-9 slices from ≥ 3-4 animals for each condition; data are plotted as mean ± SEM. Letters above bars indicate statistical groups; p < 0.05, Kruskal-Wallis test. Circles represent individual cells. (C) Representative images of immunostaining for GAD65+ puncta (red) in CA1 pyramidal cell body layer of fixed organotypic slices. Nuclei are stained for DAPI (blue). A single z-plane is shown. Due to the viruses used, neurons from the WT and Plexin-B1 KO conditions express GFP in the whole cell while neurons in the Plexin-B2 KO and double KO conditions express a Cre-GFP fusion protein which localizes to nuclei.

We found that deletion of Plexin-B1 on its own (*Plxnb1^−/−^*;*Plxnb2*^flx/flx^ + AAV.hSyn.GFP) caused a decrease in GAD65 puncta density compared to wild-type condition (*Plxnb1*^+/+^;*Plxnb2*^flx/flx^ + AAV.hSyn.GFP) (Fig. 4B,C). Interestingly, knockout of the Plexin-B2 gene on its own (*Plxnb1*^+/+^;*Plxnb2*^flx/flx^ + AAV.hSyn.Cre-eGFP) resulted in GAD65 puncta density that was significantly decreased from the GAD65 density observed in the wild-type condition, and was significantly decreased when compared to the Plexin-B1 KO condition. Deletion of both Plexin-B1 and Plexin-B2 simultaneously (*Plxnb1^−/−^*;*Plxnb2*^flx/flx^ + AAV.hSyn.Cre-eGFP) resulted in decreased GAD65 puncta density similar to Plexin-B2 knockout alone. Based on these results, we conclude that Plexin-B1 and Plexin-B2 are each required for proper GABAergic synapse formation. Our observations also suggest that the relative contribution of Plexin-B2 to GABAergic synapse development in CA1 is greater than that of Plexin-B1. Further, the fact that the decreased GAD65 puncta density observed in the double KO condition was similar to that found in the Plexin-B2 KO on its own suggests that these two genes do not function redundantly and that Plexin-B2 is epistatic to Plexin-B1.

### Engineering and validation of chimeric Plexin-B receptors

To better understand the different relative contributions of Plexin-B1 and Plexin-B2 receptors to GABAergic synapse development, we next utilized the NCBI sequence alignment tool to assess the amino acid sequence similarity between the two receptors, categorized by extracellular domain (ECD), transmembrane domain (TMD), and intracellular domain (ICD). We found that the percent conserved sequence identity between Plexin-B1 and Plexin-B2 peptides was 38%, 15%, and 61% for the ECD, TMD, and ICD, respectively (Table 1). The designation of the sequences of the ECD, TMD, and ICD are also indicated in Table 1. We then designed chimeric Plexin-B1 and Plexin-B2 receptors in which amino acid sequences of the ECD or ICD were swapped between Plexin-B1 and Plexin-B2. Recent computational predictions assert that the TMD of Plexin-B receptors may play a previously unappreciated role in signaling and gating signal transduction (Zhang et al., 2015). Thus, we aimed to isolate the function of the TMD by creating two sets of two chimeras that differed only in the sequence of the TMD (Fig. 5A). Finally, all constructs also contained an N-terminal myc tag for protein detection. To validate these constructs, we asked if the Plexin-B chimeras trafficked to the cell membrane and retained the capacity to signal to the cytoskeleton.

**Figure 5.**
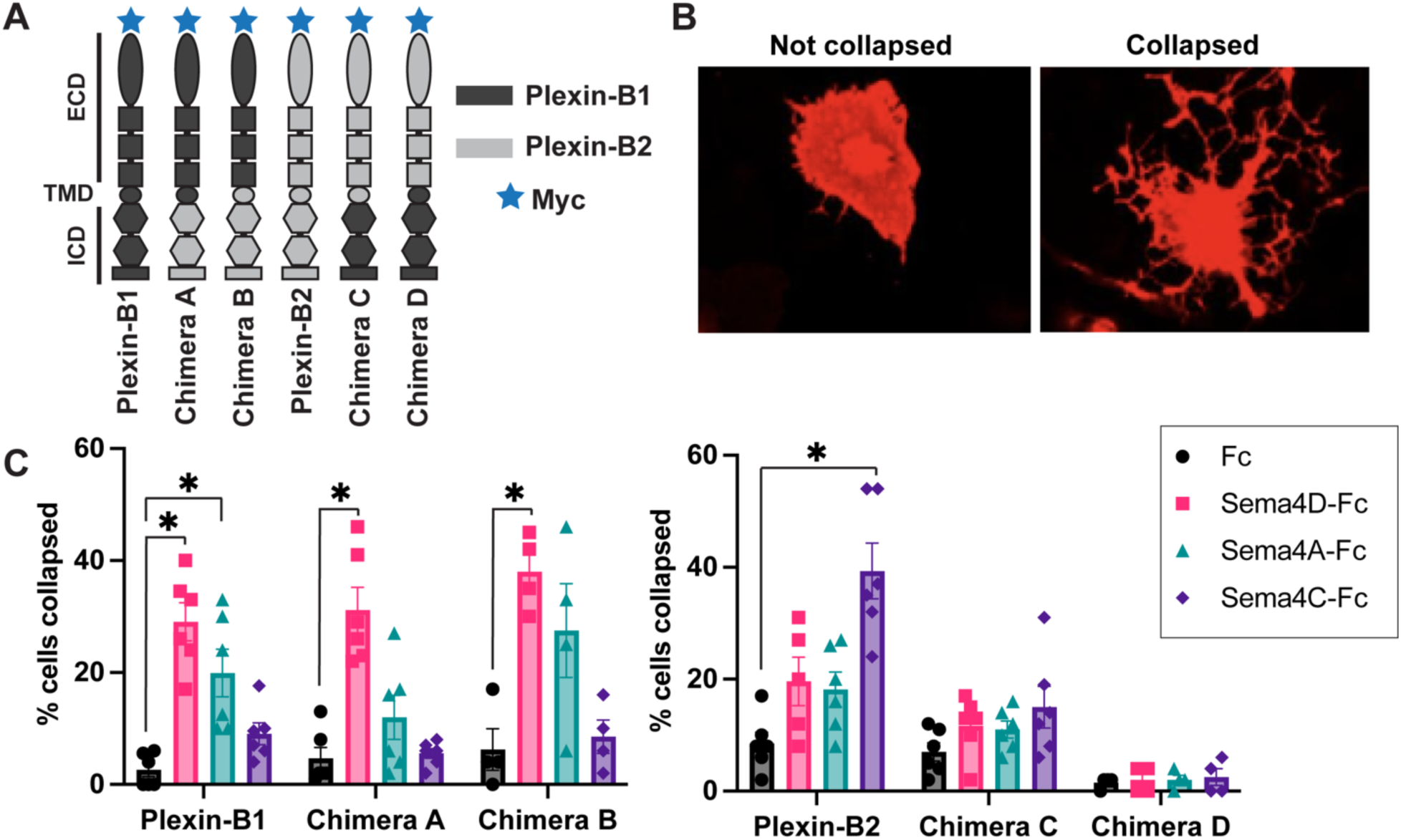
Validation of chimeric Plexin-B receptors in a COS-7 cell functional assay. (A) Schematic diagram illustrating WT Plexin-B1 and Plexin-B2 and chimeric receptors. (B) Representative images of COS-7 cells transfected with plasmids containing WT Plexin-B1 or WT Plexin-B2 cDNAs and immunostained for surface expression of the N-terminal myc tag. (C) Cytoskeleton collapse induced by treatment with soluble ECD of class 4 Semaphorins conjugated to Fc or Fc alone. Data plotted as percentage of collapsed cells divided by total cells counted (45-60 cells sampled per coverslip). n = 4-6 coverslips from 2-3 experiments per condition. For each receptor, different ligand treatment conditions were compared to the Fc control. One-Way ANOVA with Tukey’s test for multiple comparisons; * p <0.05. Points represent individual coverslips.

**Table 1.** Molecular domain designations for wild type and chimeric Plexin-B receptor constructs.

| Gene | ECD sequence | TMD sequence | ICD sequence |
| --- | --- | --- | --- |
| Plexin-B1 | NP_001123554.1<br>aa19 -1486 | NP_001123554.1<br>aa1487-1510 | NP_001123554.1<br>aa1511-2135 |
| Plexin-B2 | <u>NP_001363795.1</u><br>aa20-1197 | <u>NP_001363795.1</u><br>aa1198-1219 | <u>NP_001363795.1</u><br>aa1220-1838 |
| Amino acid<br>sequence similarity | 38% | 15% | 61% |

We transfected COS-7 cells with WT Plexin-B1, WT Plexin-B2, and all four chimeric Plexin-B receptor constructs. 48 hours later, we performed immunostaining using an antibody against the myc epitope tag on the N-terminus of these proteins using non permeabilizing cell fixation and immunostaining conditions to selectively visualize cell surface proteins (Fig. S2A). Signal for the N-terminal myc tag was observed at the cell surface for all constructs tested, indicating that WT and chimeric Plexin-B receptors are expressed and trafficked to the cell membrane (Fig. S2B).

One well-established example of Plexin-B receptor signaling to the actin cytoskeleton is observable by the cell collapse assay. In this assay, exogenous expression of a Plexin-B receptor in COS-7 cells followed by application of a high binding affinity class 4 Semaphorin ligand induces cytoskeleton collapse (Oinuma et al., 2003; Yukawa et al., 2010). Thus, we utilized this collapse assay to determine if the chimeric receptors also were able to transduce signals to the cytoskeleton upon Semaphorin ligand binding. To begin, we transfected COS-7 cells with WT Plexin-B1, WT Plexin-B2, or each of the four chimeric Plexin-B receptors. Forty-eight hours after transfection we incubated the cells for 1.5 hours with Sema4D-Fc, Sema4A-Fc, Sema4C-Fc or Fc control. Following fixation, cells were immunostained for the N-terminal myc tag to identify transfected cells and myc+ cells were classified as collapsed or not based on morphology of the cell (Fig. 5B).

As expected, the Fc control treatment did not cause collapse of COS-7 expressing Plexin-B1, Plexin-B2, or Chimeras A-D (Fig. 5C). Plexin-B2-expressing cells treated with Sema4C-Fc exhibited the greatest amount of cell collapse as compared to all other conditions. Plexin-B1-expressing cells treated with Sema4D-Fc demonstrated the second-highest amount of collapse; these observations are in line with a previous study which showed that Sema4C binds with higher affinity to Plexin-B2 than Sema4D does to Plexin-B1 (Deng et al., 2007). Cells expressing Plexin-B1, but not those expressing Plexin-B2, collapsed in response to treatment with Sema4A-Fc.

Cells expressing Chimera A or Chimera B behaved very similarly to each other and collapsed in response to Sema4D-Fc, but not in response to any other ligand (Fig. 5C), consistent with the fact that Chimera A and B encode the Plexin-B1-derived ECD. Notably, Chimeras C and D also behaved similarly to each other and did not collapse in response to treatment with any ligand, suggesting that Chimeras C and D are nonfunctional, perhaps either due to lack of ligand-binding or to protein misfolding. Based on these results we chose to focus our subsequent experiments on Chimeras A and B.

### Chimeric Plexin-B receptors fail to instruct proper GABAergic synapse formation

We next asked whether Chimera A or Chimera B is sufficient to rescue the deficit in GABAergic synapse formation caused by deletion of Plexin-B2. To do this, we packaged WT Plexin-B2, Chimera A, or Chimera B into adenovirus downstream of a FLEX cassette in order to drive Cre-dependent expression of these genes. To assay GABAergic synapse density, dissociated hippocampal cultures were isolated from P0-P1 pups (*Plxnb2*^flx/flx^) and infected with a combination of appropriate viruses at day 5-6 post infection which serve the following purposes (Table 2): AAV9.hSyn.GFP fills the cell with GFP. AAV9.hSyn.Cre-P2A-TdTomato conditionally deletes Plexin-B2 while filling the cell with TdTomato, and ad.FLEX.Plexin-B2, ad.FLEX.Chimera-A, or ad.FLEX.Chimera-B drive Cre-dependent expression of WT Plexin-B2 or chimeric Plexin-B receptors (Table 2). Note that due to the high infection efficiency of these viruses (approximately 80% of cells infected, data not shown), both the pre and postsynaptic neurons are expressing these proteins. At DIV16, we fixed and stained the cultures using antibodies that recognize GAD65 and gephyrin, a scaffolding protein localized to the postsynaptic specialization of GABAergic synapses, followed by confocal imaging. We analyzed the density of GAD65 and/or gephyrin puncta onto cell somas that were either GFP+ or TdTomato+, demonstrating their successful infection with indicated viruses.

**Table 2.** Infection conditions used in dissociated Plexin-B2 cKO dissociated neuronal cultures.

| Condition | Genotype | Virus 1 | Virus 2 | Cre recombinase delivered? | Other genes delivered |
| --- | --- | --- | --- | --- | --- |
| **Control** | *Plxnb2<sup>flx/flx</sup>* | AAV.hSyn.GFP | None | No | None |
| **cKO** | *Plxnb2<sup>flx/flx</sup>* | AAV.hSyn.Cre-P2A-TdTomato | None | Yes | None |
| **cKO + Plexin-B2** | *Plxnb2<sup>flx/flx</sup>* | AAV.hSyn.Cre-P2A-TdTomato | ad.FLEX.Plexin-B2 | Yes | Plexin-B2 |
| **cKO + Chimera A** | *Plxnb2<sup>flx/flx</sup>* | AAV.hSyn.Cre-P2A-TdTomato | ad.FLEX.Chimera-A | Yes | Chimera A |
| **cKO + Chimera B** | *Plxnb2<sup>flx/flx</sup>* | AAV.hSyn.Cre-P2A-TdTomato | ad.FLEX.Chimera-B | Yes | Chimera B |

We found that the density of GABAergic synapses, defined as the overlap of GAD65 and gephyrin puncta onto a TdTomato- or GFP-expressing neuronal soma, was significantly reduced in the absence of Plexin-B2 (Fig. 6A-C). This deficit was partially restored by expression of WT Plexin-B2; however, neither Chimera A nor Chimera B restored the decreased density of GABAergic synapses observed upon Plexin-B2 cKO. We also found that expression of Plexin-B1 did not restore proper GABAergic synapse development in the absence of Plexin-B2 (Fig. S3A). Further, we examined GABAergic synapse density in the proximal dendrites of the hippocampal pyramidal postsynaptic neuron and found no change upon Plexin-B2 cKO (Fig. S4A-D).

**Figure 6.**
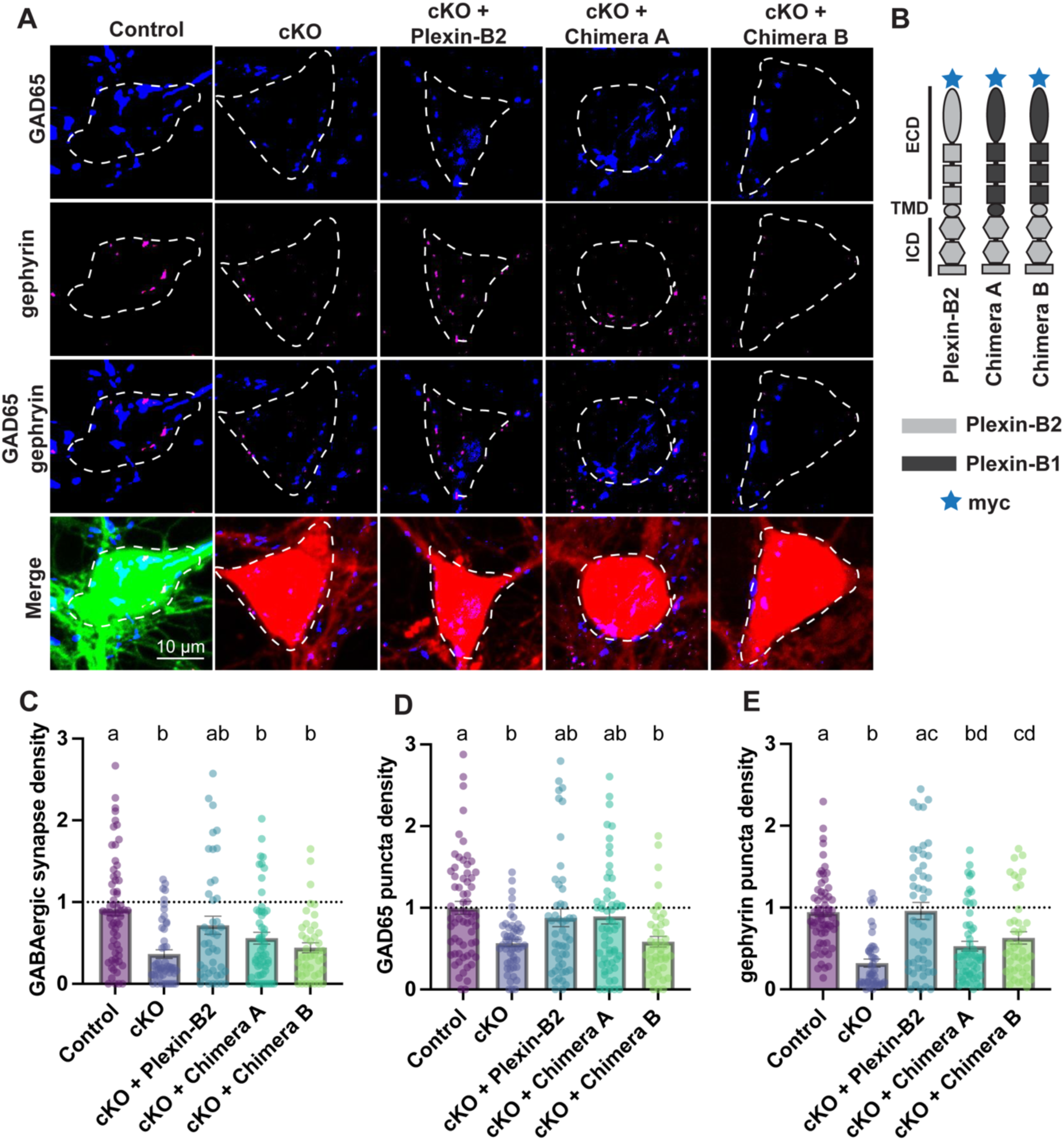
Chimeric Plexin-B receptors are not sufficient to restore GABAergic synapse formation in the absence of Plexin-B2. Neurons from *Plxnb2^flx/flx^* mice were infected at DIV5/6 with viruses indicated in Table 2. At DIV16, neurons were fixed and immunostained for the indicated synaptic markers. (A) Representative images of soma from each condition immunostained for GAD65 (blue) and gephyrin (magenta). Dotted white line shows the soma boundary. Panels labeled “merge” show GAD65, gephyrin, as well as the cell fill (either GFP or TdTomato). (B) Schematic representation of WT and chimeric Plexin-B receptors delivered to neurons. (C) Density of GABAergic synapses, defined by the overlap of GAD65/gephyrin puncta on a GFP- or TdTomato-labeled soma, divided by area of the soma. (D) Density of GAD65-positive puncta divided by area of the soma. (E) Density of gephyrin-positive puncta divided by the area of the soma. (C-E) Data are normalized to the control condition and outliers were removed (see methods). n ≥ 45 neurons per condition from at least three experiments; p ≤ 0.05, Kruskal-Wallis test. Statistical differences are indicated by different letters and bars sharing a letter are not statistically different. Error bars are ± standard error of the average ratio. Note: the control condition in some cases is < 1 due to outlier removal; see methods. Scale bar, 10 µm. Circles represent values for individual cells.

We next assessed the effect of expression of each chimera on formation of the presynaptic GABAergic bouton (as indicated by GAD65 staining alone) or the postsynaptic specialization (as indicated by gephyrin staining alone). To this end, we quantified the density of GAD65 puncta localized to cell somas or gephyrin puncta localized to cell somas without demanding co-localization between the GAD65/gephyrin signals. We found that the density of both GAD65 and gephyrin puncta on the cell soma was significantly decreased by Plexin-B2 cKO and that this deficit was partially restored by expression of WT Plexin-B2 (Fig. 6D, E). Thus, neuronal Plexin-B2 expression is required for proper formation of both the inhibitory pre- and postsynaptic specializations. However, because Plexin-B2 is knocked out in both the pre- and postsynaptic neurons, we do not know if this reflects a separate role for Plexin-B2 in each neuron or a trans-synaptic signaling complex. Surprisingly, expression of Chimera A in the Plexin-B2 knockout neurons resulted in partial restoration of the presynaptic specialization formation, while expression of Chimera B failed to restore presynaptic specialization formation (Fig. 6D). Chimera A and B differ only in the identity of the TMD; thus, these observations indicate a functional distinction between the TMD of Plexin-B1 and Plexin-B2. Further, expression of Plexin-B1 in Plexin-B2 knockout neurons failed to restore assembly of the presynaptic specialization (Fig. S3B). Comparison of Plexin-B1 with Chimera A and Chimera B indicates that the Plexin-B2 ICD plays a critical role in formation of the presynaptic specialization, downstream of a process that can be engaged by the TMD of Plexin-B1 but not of Plexin-B2.

Next, examination of the postsynaptic marker on its own (gephyrin) revealed that expression of neither Chimera A nor Chimera B in Plexin-B2 knockout neurons was able to drive proper formation of the postsynaptic specialization. Notably, expression of Chimera B resulted in a density of gephyrin puncta mid-way between the density of gephyrin puncta in Plexin-B2 cKO and WT conditions, indicating a partial restoration of formation of the postsynaptic specialization (Fig. 6E). Chimera A and B both lack the Plexin-B2 ECD. Thus, our observations taken together indicate that the Plexin-B2 ECD is required for formation of the GABAergic postsynaptic, but not presynaptic specialization. Finally, expression of Plexin-B1 in Plexin-B2 knockout neurons partially restored formation of the postsynaptic specialization (p=0.08 compared to control but not statistically different from the Plexin-B2 cKO condition; p>0.05) (Fig. S3C).

### Glutamatergic synapse density is unaffected by Plexin-B2 knockout

We next interrogated a role for Plexin-B2 in glutamatergic synapse development. Again, at DIV5/DIV6 we infected dissociated hippocampal cultures from *Plxnb2*^flx/flx^ mice with viruses as described above and in Table 2. At DIV16, we fixed and stained the cultures using antibodies that recognize VGLUT1, a presynaptic component of glutamatergic synapses. We analyzed the formation of VGLUT1 puncta onto dendrites that were either GFP+ or TdTomato+. As expected, we did not observe a difference in density of VGLUT1 puncta between the WT (*Plxnb2*^flx/flx^ + AAV.hSyn.GFP) and Plexin-B2 KO (*Plxnb2*^flx/flx^ + AAV.hSyn.Cre-P2A-TdTomato) conditions (Fig. 7A, B). Driving expression of WT Plexin-B2, Chimera A, or Chimera B, in the Plexin-B2 KO background also had no effect on glutamatergic synapse density. Thus, we conclude that Plexin-B2 is not required for proper formation of glutamatergic synapses.

**Figure 7.**
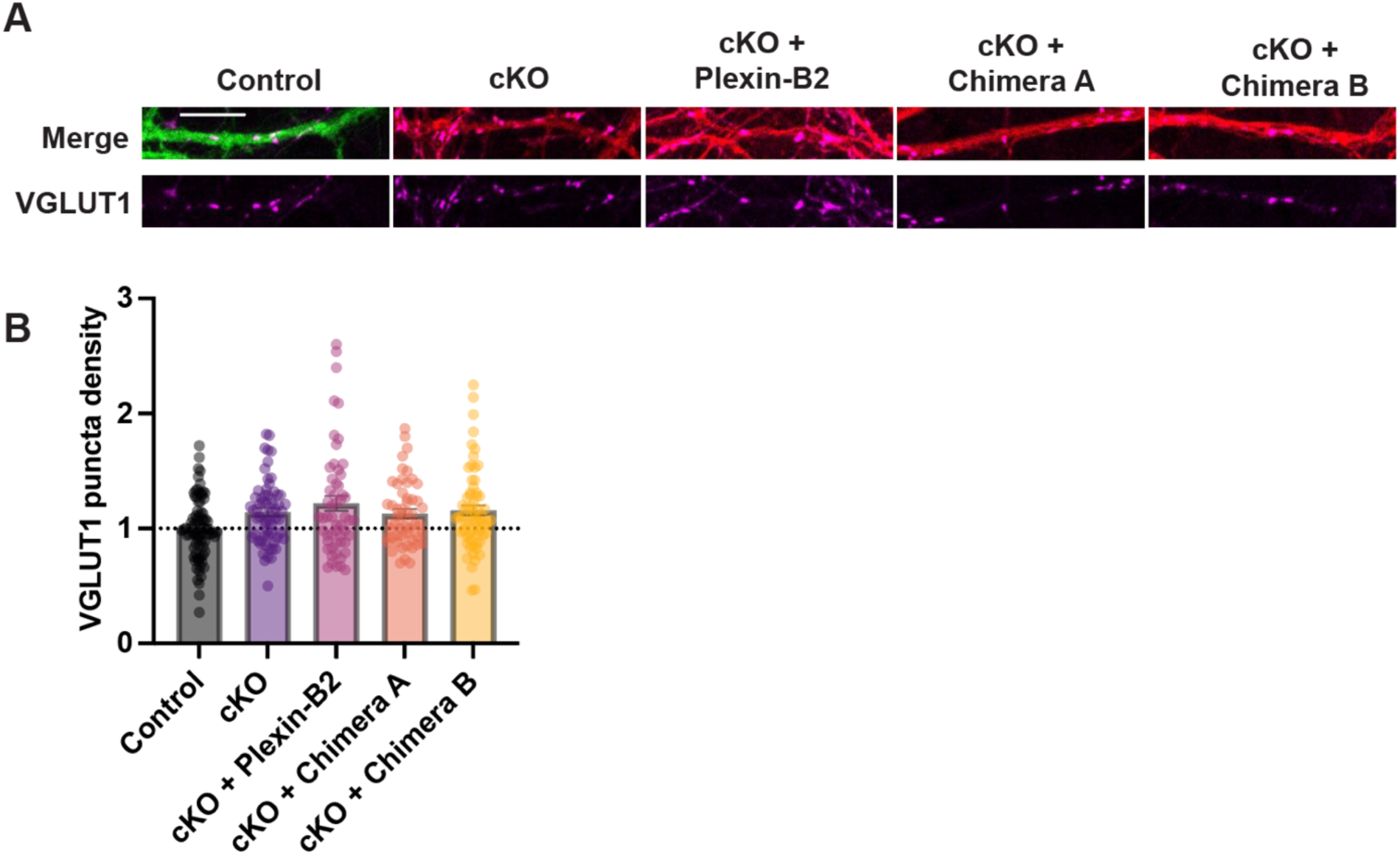
Plexin-B2 does not regulate glutamatergic synapse formation. (A) Representative dendrite stretches from each condition immunostained for VGLUT1 (magenta). Cell fill is either GFP or TdTomato. (B) Quantification of the number of VGLUT1-positive puncta on proximal dendrites in a given field of view, divided by the area of the dendrites. Data are normalized to the average density of the GFP control condition and outliers were removed (see methods). n ≥ 45 neurons per condition from at least three experiments; no significant differences between conditions were found according to a Kruskal-Wallis test. Statistical differences are indicated by different letters and bars sharing a letter are not statistically different. Error bars are ± standard error of the average ratio. Scale bars, 10 μm. Circles represent values for individual fields of view.

## Methods

### Ethics statement

All animal procedures were approved by the Brandeis University Institutional Animal Care and Usage Committee, and all experiments were performed in accordance with relevant guidelines and regulations.

### Animals

To obtain PV^Cre^;Plxnb2^+/flx^ animals, we crossed PV^Cre^ mice (Jackson Laboratories, #008069) with mice harboring a Plexin-B2^flx^ conditional allele (Plxnb2^tm1c(EUCOMM)Wtsi^) (Daviaud et al., 2016) which contain loxP sites flanking exons 7-9 of the Plxnb2 gene. Offspring of genotype PV^Cre^;Plxnb2^+/flx^ mice were crossed to each other to produce pups with the genotypes PV^Cre^;Plxnb2^+/+^, PV^Cre^;Plxnb2^+/flx^, and PV^Cre^;Plxnb2^flx/flx^. All mice used in this study were on the C57BL/6 background.

### In situ hybridization

Wild type C57BL/6 mouse brains (P14) were rapidly dissected over ice and covered in OCT (TissueTek) in plastic tissue molds. Blocks were flash frozen in 2-methylbutane chilled in a liquid nitrogen bath and stored at −80°C for up to 2 weeks. Tissue blocks were warmed to −20°C for at least 1h prior to sectioning and sectioned at 18 µm using a cryostat (Leica CM3050S) at −20°C. Coronal sections were mounted on room-temperature charged glass slides (Colorfrost Plus, Fisher), stored at −20°C until sectioning was completed and transferred to a −80°C freezer for up to 3 months. Mounted sections were fixed in 4% paraformaldehyde in PBS and dehydrated by successive washes with 50%, 70%, and 100% EtOH. Dried slides were pretreated with RNAScope hydrogen peroxide reagent (Advanced Cell Diagnostics) for 10 minutes at RT and RNAscope Protease Plus for 30 minutes at RT. Probes (see Table 3 below) were then hybridized for 2h at 40°C in a humidified hybridization oven. Slides were stored overnight at RT in 5x SSC. Probe amplification and detection steps were followed according to manufacturer directions for RNAscope 2.5 HD Duplex kit. Following the final amplification step, nuclei were counterstained with 50% Gill’s Hematoxylin I solution, briefly rinsed with tap water, and treated with bluing reagent (0.02% ammonia) for 10s. Slides were then baked in a 60°C dry oven for 20 minutes, dipped in xylene, and mounted with Vectamount permanent mounting solution.

**Table 3.** Probes used for *in situ* hybridization experiments.

| Reagent | Cat # | Supplier |
| --- | --- | --- |
| RNAscope® 2.5 HD Duplex Reagent Kit | 322430 | Advanced Cell<br>Diagnostics |
| RNAscope® 2.5 Duplex Positive Control probe - Mm | 321651 | Advanced Cell<br>Diagnostics |
| RNAscope® 2-plex Negative Control Probe | 320751 | Advanced Cell<br>Diagnostics |
| RNAscope Probe - Mm-Pvalb-C2 | 421931-C2 | Advanced Cell<br>Diagnostics |
| RNAscope Probe - Mm-Plxnb2 | 459181 | Advanced Cell<br>Diagnostics |

### Organotypic slice culture

Plxnb2^flx/flx^ mice of both sexes were sacrificed between P6-P8 and their brains were harvested into cutting solution (126 mM NaCl, 25 mM NaHCO3, 3 mM KCl, 1 mM NaH2PO4, 20 mM dextrose, 2 mM CaCl2, 2 mM MgCl2 in deionized water at 315–319 mOsm) (adapted from Stoppini et al., 1991)). Coronal slices were taken at a thickness of 300 µm using a tissue chopper (Compresstome VF-200, Precisionary Instruments Inc.). Individual slices were placed on cell culture inserts (0.4 µm pore size, Millipore). Organotypic culture media (2 mM glutamax, 1 mM CaCl2, 2 mM MgSO4, 12.9 mM d-glucose, 0.08% ascorbic acid, 18 mM NaHCO3, 35 mM HEPES, 20% horse serum, 1 mg/mL insulin in minimum essential media) at pH 7.45 and 305 mOsm was added outside of the inserts. Slices were maintained for 6 days in vitro at 35 °C and 5% CO2 with media replacements every other day.

One day after harvesting slices, a solution containing either AAV9.hSyn.eGFP.WPRE.bGH or AAV9.hSyn.HI.eGFP-Cre.WPRE.SV40 (Addgene; #50465-AAV9 or #105540-AAV9) was pipetted onto the hippocampus within the slice (1 μL on each hemisphere, each virus at 1.8 ×10^12^ Gc/mL). At DIV6, slices were fixed in 4% paraformaldehyde/4% sucrose for 20 min at 4 °C.

Images were obtained using a Nikon Eclipse Ti-E inverted motorized microscope equipped with a Swiftcam 16 Megapixel RGB sideport camera at either 10x or 60x magnification. Single-plane full spectrum brightfield images were obtained under white light using built-in Swift Imaging 3.0 software. To avoid potential bias in image acquisition, all visible Pvalb+ cells within the hippocampal CA1, CA2, CA3, and DG regions were captured. Within each experiment, images were acquired with identical settings for illumination brightness, gain, and exposure.

Duplex images were analyzed using QuPath 0.3.2 software. Automated detection of hematoxylin-counterstained nuclei was performed with a background radius of 24 µm, sigma factor of 1.8 µm, and quality threshold of 0.08. Next, the subcellular spot detection feature was used to quantify puncta within cell nuclei. Detection was performed with an expected spot size of 2 µm and a range of 0.5-4 µm. Detection thresholds for each marker were determined empirically. After detection of spots and clusters (a cluster is a detected feature that exceeds the expected spot size), a cell was classified as Pvalb+ if it contained at least one Pvalb cluster and nuclear optical density of the Pvalb marker exceeded a set threshold (optical density/nuclear area ratio > 1). As Pvalb mRNA transcript levels are very high Pvalb+ cells, this threshold reliably ensured that only cells unambiguously expressing Pvalb were included for analysis. A cell was classified as expressing Pvalb or Plxnb2 if it contained at least one mRNA cluster or at least 4 spots, as previously described (McDermott et al., 2018).

### Intracerebroventricular Injection

Intracerebroventricular (ICV) viral injections were performed on PV^Cre^;Plxnb2^flx/flx^ pups aged P1-3 (adapted from Kim et al., 2014). Mice were injected ⅓ of the distance between the eye of the pup and lambda at a depth of ∼3 mm. 2 ul per hemisphere of AAV.FLEX.TdTomato at a concentration of 1.5 x 10^12^ Gc/mL was delivered using a custom Hamilton syringe with the following custom specifications: 32GA, 1 in length, 12 degrees, point style 4. At ∼3 months of age, mice were transcardially perfused with sterile artificial cerebrospinal fluid followed by 4% paraformaldehyde with 4% sucrose.

Brains dissected out and kept in the paraformaldehyde solution at 4°C overnight, after which point they were washed 3 times for 10 minutes with 1x PBS. Following, brains were embedded in a 2% agar solution and sliced on a Leica VT1000 S vibratome at a section thickness of 45 µM. Slices containing the medial hippocampus were prepared for immunostaining.

### COS-7 cell culture assays

#### Collapse assays

COS-7 were grown as above and seeded at a density of 15,000/well directly onto sterile plastic 24-well plates. 24 h afterwards, COS-7 cells were transiently transfected with relevant cDNAs using a lipofectamine 3000 kit (Thermofisher). 48 h later COS-7 cells were incubated with the soluble extracellular domains of Sema4D, Sema4A, or Sema4C, conjugated to Fc (R&D Systems, #7470-S4, 4694-S4, 6125-S4) or with Fc (R&D Systems, #110-HG) for 1.5 hours at 37°C prior to fixation. Following, immunostaining for myc was performed. Coverslips were mounted on glass slides using Aquamount and manually scored for collapse behavior using a 20x objective on a BX-Z 710 Fluorescence Microscope (Keyence).

### Neuronal cell culture

First, primary mouse glia were plated onto 12 mm glass coverslips that had been coated overnight with poly-d-lysine (20 μg/ml) and laminin (3.4 μg/ml) in 24 well plates. Just before plating glia, coverslips were washed twice with deionized water and then once with DMEM. When glia formed a confluent feeder layer, hippocampi were harvested from Plxb2^flx/flx^ pups aged P1-P2 of both sexes, dissociated, and plated directly onto the glia. Cultured cells were fed every 3-4 days by replacing half the media per well with fresh media with AraC. Tail samples from all pups within a single experiment were combined and the genotype was confirmed by PCR to detect the wild type or floxed version of the Plxnb2 allele.

For synapse assays, neurons were infected on DIV4 or DIV5 with either AAV.hSyn.GFP (Addgene; #50465-AAV9) or AAV.hSyn-Cre-P2A-dTomato (Addgene; #107738-AAV9) at a final concentration of 4×10^9^ Gc/mL, or with a combination of AAV.hSyn-Cre-P2A-dTomato plus 1 ul one of the following at a final concentration of 1×10^6^ PFU: ad.FLEX.Plexin-B2, ad.FLEX.Chimera-A, or ad.FLEX.Chimera-B. Cultures were maintained with half media changes every 3-4 days until being fixed with 4% paraformaldehyde for 8 minutes at DIV16.

### Immunostaining

All antibody information including final working dilutions used are indicated in Table 4.

**Table 4.** Antibody information.

| Antibody | Company | Cat number | Related to experiments in Figure | Dilution |
| --- | --- | --- | --- | --- |
| Goat-anti-chicken Alexa Fluor 405 | Abcam | ab175674 | Figures 3 & 4 | 1:500 |
| Chicken-anti-PV | Synaptic Systems | 195 006 | Figure 3 & 4 | 1:300 |
| Mouse-anti-GAD65 | MilliporeSigma | MAB351 | Figure 3, 4, 6 & Supp. Figure 4 | 1:300 |
| Alexa Fluor 488 goat-anti-mouse IgG (H+L) | Invitrogen | A11029 | Figure 3 & 4 | 1:500 |
| Alexa Fluor 633 goat anti-chicken IgG (H+L) | Invitrogen | A21103 | Figure 3 & 4 | 1:500 |
| Mouse-anti-myc (9E10) | Santa Cruz | Sc-40 | Figure 5 and Supp. Figure 2 | 1:500 (permeabilized) or 1:50 (surface) |
| Donkey-anti-mouse Alexa Fluor 405 | Abcam | ab175659 | Figure 6 | 1:1000 |
| Alexa Fluor 633 goat-anti-guinea pig IgG (H+L) | Invitrogen | A21105 | Figure 6 | 1:1000 |
| Guinea pig-anti-gephyrin | Synaptic Systems | 147 318 | Figure 6 | 1:1000 |
| Guinea pig-anti-VGLUT1 | Synaptic Systems | 135 304 | Figure 7 | 1:1000 |
| Mouse-anti-TFIIB | Santa Cruz Biotechnology | sc-271736 | Supp. Figure 2 | 1:50 (surface) or 1:500 (permeabilized) |

#### Organotypic slices

After fixation, slices underwent 3 × 10-minute washes in PBS and were incubated overnight in permeabilization solution (0.5% Triton-X in PBS) followed by another overnight incubation in blocking solution (20% bovine serum albumin with 0.1% Triton-X in PBS) at 4°C. The next day, slices were incubated with primary antibodies diluted in blocking solution and incubated overnight at 4°C. Following, slices were washed 3 × 10-minutes in PBS and incubated for 2hrs at room temperature with fluorescently conjugated secondary antibodies in secondary solution (blocking solution diluted 1:1000 in PBS). After final 3 × 10-minute PBS washes, organotypic slices were mounted (tissue side up) on slides in Vectashield + DAPI mounting media (Vector Laboratories).

Perfused tissue sections from PV^Cre^;Plxnb2^+/+^ or PV^cre^;Plxnb2^flx/flx^ mice: These slices underwent the same immunostaining protocol as the organotypic slices with the exception of which antibodies were applied and at which working dilutions. Also, these sections were mounted in Aquamount, not Vectashield + DAPI.

#### COS-7 cells and dissociated neurons

Cells were fixed with 4% paraformaldehyde/4% sucrose for 8 min at room temperature. Coverslips were washed 3 times with 1× PBS for 5 min each and incubated overnight at 4°C with blocking solution (3% BSA plus 0.1% Triton X) and the next day with primary antibodies also overnight at 4°C. All antibody dilutions were prepared in blocking solution. After overnight incubation, coverslips were washed 3 times with 1× PBS for 10 min each and then incubated with secondary antibodies (1:1000 each) in diluted blocking buffer (1:1000) for 2 h at room temperature. Coverslips were then washed 3 times with 1× PBS for 10 min each, dipped in dH2O, and mounted on glass slides with Aquamount (VWR). Where indicated, DAPI (1:5000) was added to the last PBS wash of coverslips containing COS-7 cells.

### DNA plasmids and viruses

Human Plexin-B1 and Plexin-B2 full length cDNA were cloned from pcDNA2-vsv-Plexin-B1 and pMT2-Plexin-B2-vsv (gifts from Dr. Luca Tamagnone) into a mammalian expression vector containing a CMV promoter, artificial signal sequence, and myc epitope tag at the N-terminus, resulting in pcmv-myc-Plexin-B1 and pcmv-myc-Plexin-B1 constructs. Chimeric Plexin-B constructs containing Plexin-B1 and Plexin-B2 domain swaps were produced by Genscript. Target sequences for the ECD, TMD, and ICD were determined by their annotations on NCBI and are described in Table 1. To generate viruses, myc-Plexin-B2, myc-Chimera-A, and myc-Chimera-B were cloned and packaged into adenovirus. Vector Biolabs (Malvern, PA) carried out cloning, viral packaging, and purification.

### Image acquisition and data analysis

Images were acquired and analyzed by an experimenter blinded to the condition. Microscope settings for laser power, detector gain, and amplifier offset were initially optimized across multiple control coverslips to avoid oversaturation of pixels and were kept constant within each experimental round. All image analyses were conducted using ImageJ software (NIH). Specific information for each experiment is detailed below.

Synapse assays in tissue from PV^Cre^ mice: Images were acquired on a Zeiss LSM880 confocal microscope equipped with Airy scan technology using the Plan-Apochromat 63x/1.40 Oil DIC M27 objective. We acquired z-stacks (0.5 µm step size) of all cell somae satisfying the following criteria: TdTomato+/PV+, and located within the pyramidal cell layer or stratum oriens region of the CA1, CA2, or CA3 subregions of the hippocampus. For analysis of GABAergic synapse formation onto PV neurons, a single z-plane containing a PV/TdTomato double positive cell soma was chosen for each cell, an ROI was drawn around the cell soma using the TdTomato signal as a guide and the area was measured. A uniform threshold was defined for the GAD65 signal across all images and then the number of GAD65+ puncta was measured using the “Analyze Particles” function according to the same parameters used in the organotypic slices. Number of GAD65+ puncta per area was calculated and this was defined as the GAD65 puncta density. For analysis of GABAergic synapse density onto pyramidal neurons, the same optical section and methods were utilized; however, in this case an approximately rectangular ROI was drawn around an area of CA1 pyramidal cell soma layer that did not contain a PV positive soma. GAD65 puncta density was again defined as the number of GAD65+ puncta divided by the ROI area.

#### Organotypic slices

16-bit images of neurons were acquired on a Zeiss LSM880 Confocal microscope using a Plan-Apochromat 63x/1.40 Oil DIC M27 objective. Images were acquired as z-stacks (5–15 optical sections, 0.5 µm step size) for each of 4–5 fields of view per hemisphere (134.95 µm x 134.95 µm) containing the pyramidal cell layer in CA1 from each slice. Within each z-stack, the image with the greatest fluorescent intensity for GAD65 signal was chosen for analysis; neurons were selected at random for analysis using only the GFP signal, thus blinding the experimenter to the GAD65 signal in the chosen neurons. Using the DAPI signal as a guide, the nuclei of these cells were traced and then a concentric perimeter expanding a uniform distance of 2 µm around the perimeter of the nucleus was drawn using the “draw band” function in ImageJ (NIH). This defines a doughnut-shaped region of interest (ROI) around the outer perimeter of the nucleus and provides a close approximation of the cell soma boundary encompassing the area where synapses form onto the cell soma. Using ImageJ, the background signal was measured in three distinct areas of each image that lacked synaptic signal and the average mean background intensity was subtracted from the entire image. Finally, signal in the GAD65 channel was binarized, the adjustable watershed algorithm was applied, and GAD65 puncta within each ROI were quantified using the “analyze particles” function (size=0.1–10 and circularity=0.01–1.00). The number of puncta per ROI was divided by the ROI area to give a per soma density of GAD65 puncta. Between 16 and 40 cells were analyzed per slice. A subset of neurons in the CA1 region exhibit very high GAD65 immunofluorescence which fills the cell soma and are presumably interneurons. Such cells were excluded from analysis.

#### Synapse assays in cultured neurons

For neuron assays, each field of view acquired was centered on a GFP+ or TdTomato+ neuronal cell soma, capturing the entire soma and proximal dendrites. 16-bit images were acquired on a Zeiss LSM 880 confocal microscope using a 63x oil immersion objective. Z-stacks encompassing the entire cell body were acquired for each cell with a 0.5 µm step size. After image acquisition, maximum intensity projections were generated and utilized in the analysis that followed. A GABAergic synapse was defined as the triple overlap of at least one pixel of the GFP or TdTomato filled dendrites, presynaptic antibody marker, and the postsynaptic antibody marker using the ‘image calculator’ function. For each experiment, a threshold for the channel containing each synaptic marker was determined and applied uniformly across all images. The GFP or TdTomato threshold for each image was determined independently to optimize visualization of the soma and dendrites morphology. A region of interest was manually drawn around the soma using the GFP fill, the area of which was measured prior to deleting the soma from the image. Next the GFP or TdTomato was used to create a binary mask of the dendrites by the Renyi entropy thresholding method. Synapse density was calculated by the number of puncta positive for synaptic markers, divided by the area of the region analyzed (soma or dendrites).

### Statistical analysis

Statistical analyses were performed using Prism. For each experiment, specific statistical tests are described in the text and/or figure legends. Data were tested for normality before appropriate statistical tests were applied. Where indicated, outliers were identified by the ROUT method with Q = 1% and removed from the data set.

## Discussion

In this study we demonstrated that Plexin-B1 and Plexin-B2 function non-redundantly in GABAergic synapse development and furthered our understanding of the precise signaling configurations by which Plexin-B1 and Plexin-B2 modulate assembly of both the pre- and postsynaptic specialization of GABAergic synapses. We revealed a presynaptic role for Plexin-B2 in the PV+ neurons in GABAergic synapse development between PV+ interneurons and pyramidal neurons in the hippocampus. Finally, we also presented evidence that the TMD in part underlies the divergent functions observed between Plexin-B1 and Plexin-B2 in GABAergic synapse formation, and that the Plexin-B2 ECD is critical for assembly of the GABAergic postsynaptic specialization.

Synapse formation is a multistep process which requires reorganization of the actin cytoskeleton in addition to accumulation of synaptic machinery at the membrane in both the pre- and postsynaptic neuron (reviewed in Sudhof, 2018). In this study we found that Chimeric Plexin-B receptors, Chimeras A and B, were able to signaling to the actin cytoskeleton but still failed to completely restore GABAergic synapse development. These findings suggest that processes by which Plexin-B receptors regulate the actin cytoskeleton (i.e.; via modulation of Ras- and Rho-family GTPases) (Oinuma et al., 2010; Saito et al., 2009; Wang et al., 2012), are likely insufficient to instruct proper synapse development and other mechanisms must be at play. For example, Plexin-B receptors may engage pathways that recruit synaptic scaffolding machinery to the membrane. In support of this idea, we have previously shown via live imaging that Sema4D initiates splitting of gephyrin clusters into smaller puncta, possibly representing recruitment of gephyrin to nascent synaptic sites (Kuzirian et al., 2013). Interestingly, our finding that Chimera A was able to instruct formation of the GABAergic presynaptic specialization but not the postsynaptic specialization suggests that the PlexinB1 TMD influences assembly of the presynaptic specialization, while the PlexinB2 ECD directs formation of the postsynaptic specialization.

In general, Plexin-B receptors dimerize upon ligand binding which causes increased GAP activity; these are generally thought to be homodimers (Koppel & Raper, 1998; Negishi et al., 2005). Intriguingly, recent computational predictions of helix oligomers indicated that the Plexin-B2 TMD does not form stable dimers while the Plexin-B1 TMD tends to self-associate (Zhang et al., 2015). Based on this, we propose a model whereby TMD dimerization of Plexin-B1 and Plexin-B2 partially explain their functional divergence in synaptogenic processes. In this model, homodimerization of the Plexin-B1 TMD, (either within Chimera A homodimers or heterodimers between Chimera A and endogenous Plexin-B1) is sufficient to drive assembly of the GABAergic presynaptic specialization when the Plexin-B2 ICD is present (Fig. 8). Further, Plexin-B1 and Plexin-B2 are also found in cells as heterodimers (Artigiani et al., 2003), raising the possibility that Plexin-B1 homodimers, Plexin-B1/B2 heterodimers, and Plexin-B2 homodimers have different signaling capacities to regulate GABAergic synapse formation (Fig. 8).

**Figure 8.**
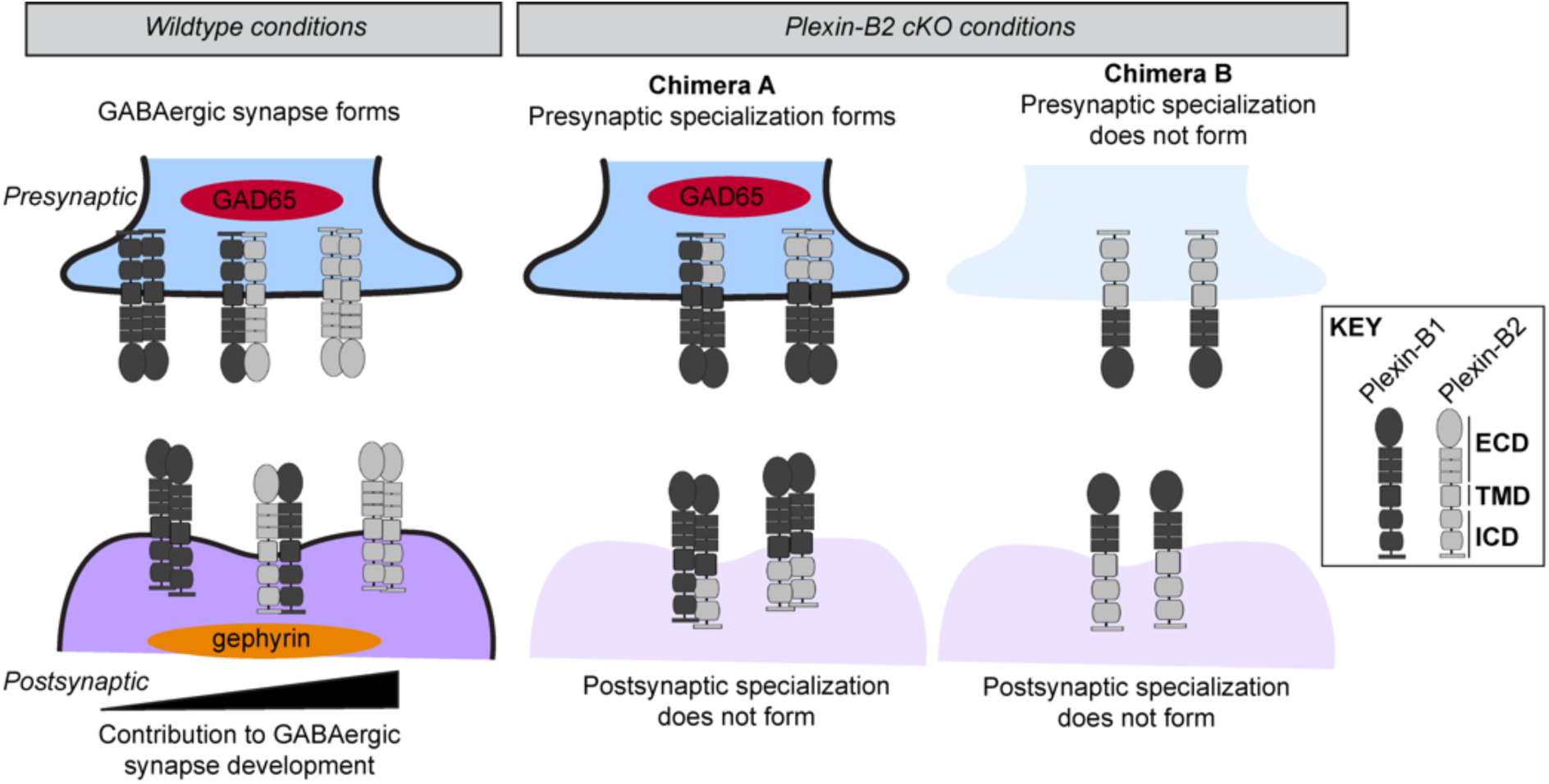
Proposed model of mechanisms of GABAergic synapse formation mediated by Plexin-B1 and Plexin-B2, and chimeric Plexin-B receptors. Middle and right panel illustrates proposed dimerization of the Plexin-B1 TMD that is sufficient to form the presynaptic specialization in the presence of the Plexin-B2 ICD, while right panel shows no dimerization and no synapse formation. Semaphorin ligands not pictured.

Our proposed model is supported by evidence that distinct and overlapping sets of receptors differentially mediate synapse formation. For example, FGFR2b drives GABAergic synapse formation while it acts synergistically with FGFR1b to drive glutamatergic synapse formation (Dabrowski et al., 2015).

Altogether our results suggest that Plexin-B TMD interactions and the dimerization state of the Plexin-B receptors may be an important mechanism regulating GABAergic synapse formation and could underlie functional mechanistic distinctions between Plexin-B1 and Plexin-B2. While the TMD has conventionally been considered simply a membrane anchor, its role in protein-protein interactions has emerged in recent years, including within synaptogenic protein families. For example, the TMD of ErbB1 and ErbB2 facilitates dimerization as well as contributes to conformational changes associated with receptor activation (Endres et al., 2013; Gerber et al., 2004). In another example, the TMD was found to contribute to homo- and heterodimerization interactions of Plexin-A/Neuropilin-1 (Aci-Seche et al., 2014). The TMD has even become a target of therapeutic interventions (Bennasroune et al., 2004; Craik et al., 2013).

Another possible mechanism underlying the distinct functions of Plexin-B1 and Plexin-B2 receptors in synapse development is that Plexin-B interaction with a coreceptor such as receptor tyrosine kinase C-MET or ErbB2 (Conrotto et al., 2004; Deng et al., 2007; Giordano et al., 2002; Swiercz et al., 2004) also contributes to this process. MET signaling is critical in Sema4D-mediated stabilization of inhibitory presynaptic boutons (Frias et al., 2019), although a role for Plexin-B1 was not assayed in this study. Differential coreceptor interactions may explain one perplexing finding from our study: Plexin-B2 deletion does not affect glutamatergic synapse density, despite previous evidence showing that Sema4A/Plexin-B2 interaction promotes glutamatergic synapse formation (McDermott et al., 2018). Thus, the role of coreceptors in Plexin-B synaptogenic signaling warrants further investigation.

Our study shows that within a given Plexin-B receptor, different molecular domains are required for assembly of either the presynaptic or postsynaptic specialization. In this study, our manipulation of Plexin-B2 expression was not constrained to either the pre- or postsynaptic neuron; thus, it is unlikely that this finding is explained by differential expression of Plexin-Bs or Chimeras in either the pre- or postsynaptic neuron. More probably, the Plexin-B2 ECD activates pathways distinct from those required for presynaptic specialization assembly. Possibilities include ECD-specific interactions with coreceptors, ECD dimerization, or possibly even through a reverse signaling mechanism by which the Plexin-B2 ECD activates a transmembrane Semaphorin either *in cis* or *in trans*. In support of the latter, one study determined that, in a reverse signaling mechanism, Plexin-B1 can serve as a receptor for Sema4A to influence cell migration (Sun et al., 2017).

An unexpected finding from this study is that Plexin-B2 KO combined with expression of exogenous full-length Plexin-B2 resulted in increased density of GABAergic synapses on dendrites of the glutamatergic neurons. One interpretation of these results is that Plexin-B2 designates the subcellular location of GABAergic synapses within the postsynaptic neuron, perhaps by being more highly localized to soma compared to dendrites. If this were to be the case, we posit that dendrites of cells expressing exogenous Plexin-B2 exhibited higher dendritic Plexin-B2 levels than in control cells, thus recruiting more inhibitory synapses to dendritic sites.

A previous study from our lab demonstrated that Plexin-B2 plays a cell-autonomous role in the formation of GABAergic synapses onto excitatory neurons (McDermott et al., 2018) but our results here demonstrate that Plexin-B2 does not play an analogous role in GABAergic synapse formation onto PV+ interneurons. However, our data also showed that Plexin-B2 is required in the PV+ neurons to form GABAergic synapses onto excitatory pyramidal neurons in the principal cell layer, suggesting that Plexin-B2 may function in both the pre- and post-synaptic neurons to regulate GABAergic synapse formation.

Our findings here have functional implications about the role of Plexin-B2 in the hippocampal circuit. PV cells are the most abundant interneuron class in the cortex and they predominantly form inputs onto the perisomatic region of pyramidal neurons; thus, PV cells tightly control excitatory transmission (Ferguson & Gao, 2018). Therefore, our findings suggest that Plexin-B2 gates inhibition in the circuit by driving formation of inhibitory PV-pyramidal cell synapses.

To our knowledge, only one other group has examined the role of Plexin-B2 in synapse development. Simonetti et al. (2021) acutely knocked out Plexin-B2 in glutamatergic neurons using a CaMKII-Cre driver line and examined gephyrin puncta density in the stratum radiatum of the CA1 region in adult mice. In contrast to our observations, Simonetti et al observed an increase in density of gephyrin puncta in stratum radiatum, which contains the apical dendrites of CA1 pyramidal cells. While the cause of the discrepancy between these two observations is not clear, possibilities include different ages of the animals used in the studies, technical differences (we analyzed gephyrin and GAD65 puncta nearer to the soma and assigned to individual cell bodies and dendrites), and the fact that our strategy knocked out Plexin-B2 in all neurons. In addition, we analyzed synapse density onto basal dendrites of hippocampal neurons, thus it could be that Plexin-B2 regulates synapse formation differently in distinct subcellular compartments, perhaps due to different innervation. In addition, the fact that apical and basal dendrites receive contacts predominantly from PV+ and CCK+ interneurons, respectively (Booker & Vida, 2018), could indicate that Plexin-B2 mediates GABAergic synapse formation between specific interneuron classes and pyramidal neurons.

In summary, our data reveal that Plexin-B1 and Plexin-B2 are functionally distinct in the context of GABAergic synapse formation and further, that different domains of the Plexin-B receptors function to promote formation of the pre- or postsynaptic specializations. Further, we uncovered an unexpected role for the TMD of Plexin-B1 receptors, suggesting that homo- and heterodimerization of Plexin-B receptors may play an important role is signaling specificity.

## ACKNOWLEDGMENTS

We thank the members of Dr. Sacha Nelson’s lab, in particular Dr. Derek Wise, for technical assistance with organotypic slice cultures and for use of equipment. We thank Dr. Lev Silberstein at the Fred Hutchinson Cancer Center for providing the *PlxnB2^flx/flx^*mice. We thank all members of the Paradis lab for insight and support. This work was supported by National Institutes of Health grants R01NS065856 (S.P.) and F31 NS118799 (S.S.A.).

