## Supplementary data for "Plexin-B1 and Plexin-B2 play non-redundant roles in GABAergic synapse formation"

### Supplementary material

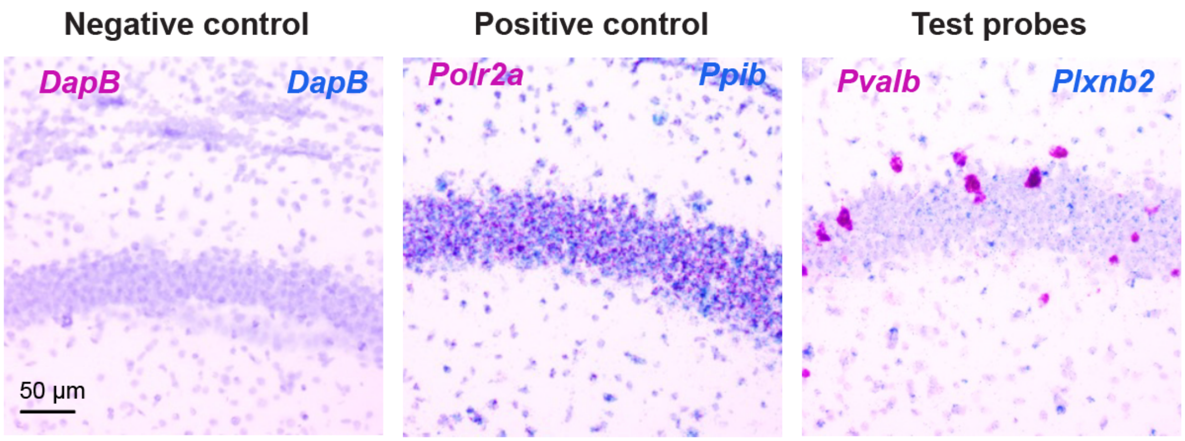


**Fig. S1. RNAscope in situ hybridization control and test probes.** Representative images of mRNA expression in the CA1 region of hippocampus in tissue isolated from P14 mice.

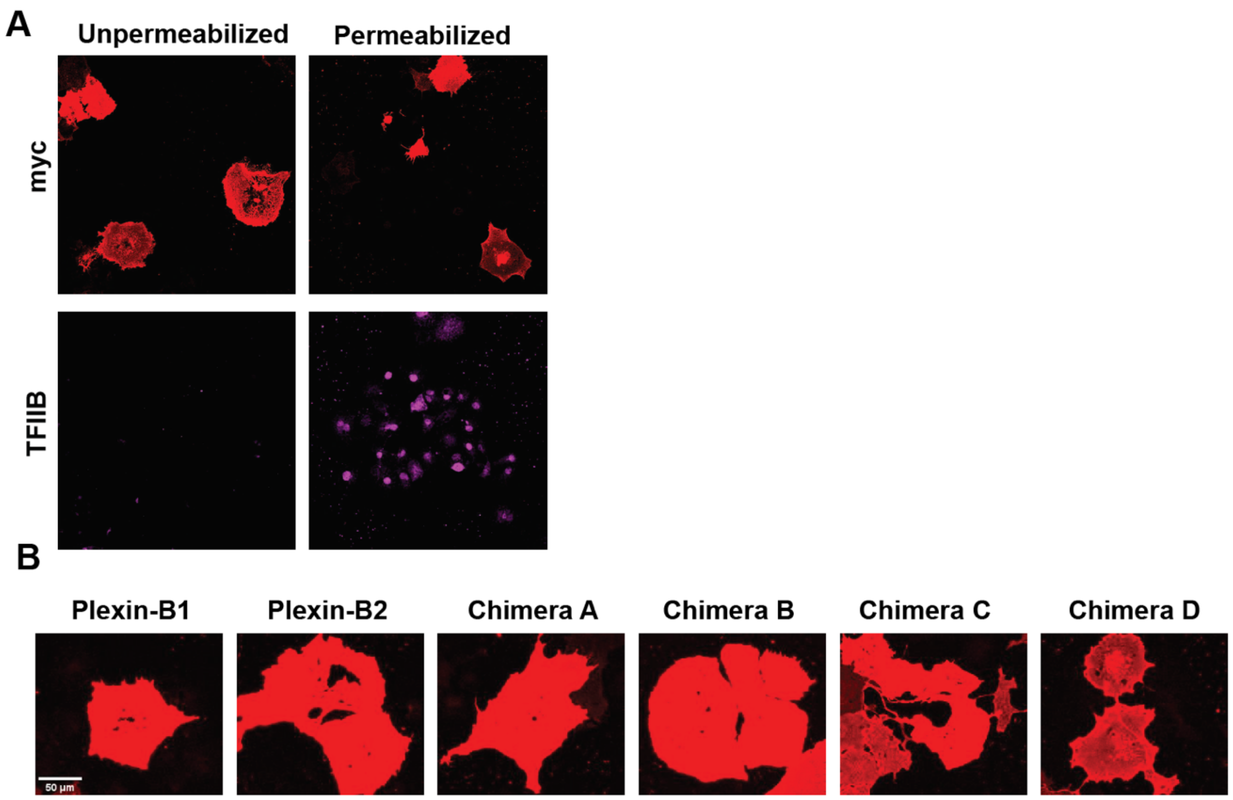


**Fig. S2. Validation of surface expression of wild type or chimeric Plexin receptors.** (A) Validation of immunostaining protocol for visualizing surface expression. COS-7 cells were transfected with a plasmid containing myc-tagged Plexin-B1. 48 hours later, cells were subjected to one of two different immunostaining protocols. To visualize Plexin-B1 localized to the cell surface, cells were fixed and stained under non-permeabilizing conditions; cells were incubated with media containing primary antibodies that recognize the myc epitope of the overexpressed Plexin-B construct or an intracellular, endogenous transcription factor (TFIIB) for 30 minutes prior to fixation (left panels). Alternatively, to visualize both intra and extracellular Plexin-B1 expression, cells were fixed, permeabilized with a detergent solution, and subsequently subjected to immunostaining using the same primary anti-myc and anti-TFIIB antibodies (right panels). Representative images showing myc or TFIIB staining are from different coverslips, as each coverslip was exposed to either anti-myc or anti-TFIIB. (B) Representative images showing cells transfected with WT or chimeric Plexin-B constructs and subjected to immunostaining for myc under non-permeabilizing conditions. Signal indicates surface expression. Scale bar shows 50 µm.

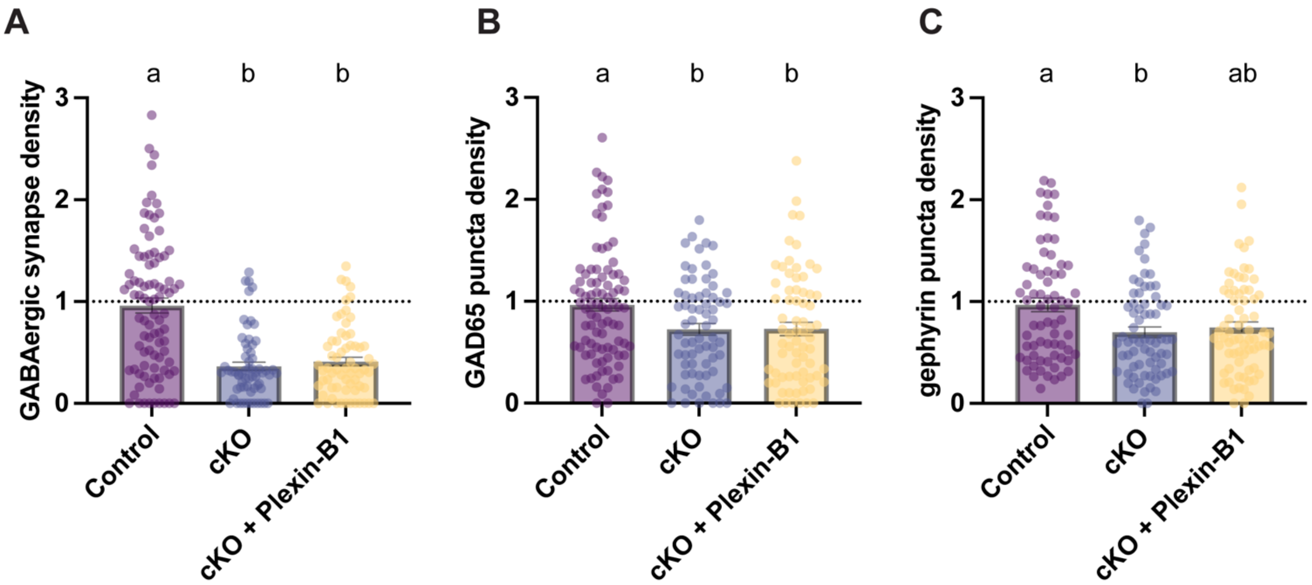


**Fig. S3. Expression of Plexin-B1 does not restore GABAergic synapse development in the absence of Plexin-B2.** (A) Representative soma each condition immunostained for GAD65 (blue) and gephyrin (magenta). (B-D) Plots showing the average density of GABAergic synapses, GAD65+ puncta, or gephyrin+ puncta on soma of individual neurons, normalized to the average density of the control condition; outliers were removed (see methods). n = 63-70 neurons per condition from three experiments; $p < 0.05$, Kruskal-Wallis test. Statistical differences are indicated by different letters and bars sharing a letter are not statistically different. Error bars are ± standard error of the mean.

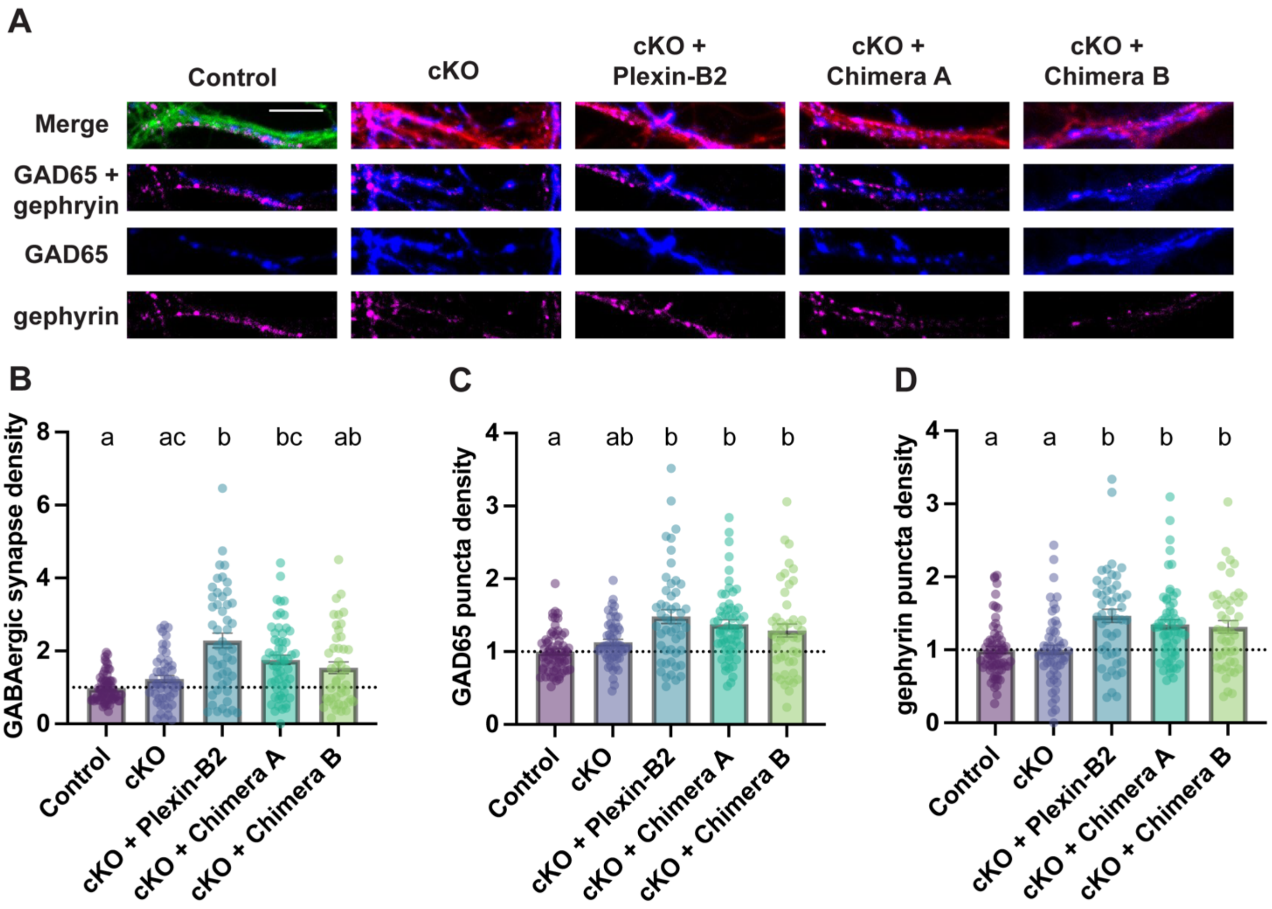


**Fig. S4. Plexin-B2 does not regulate development of GABAergic synapses onto proximal dendrites.** Neurons from *Plxnb2*$^{flx/flx}$ mice were infected at DIV5 or DIV6 with viruses indicated in Table 2. (A) Representative dendrite stretches from each condition immunostained for GAD65 (blue) and gephyrin (magenta). (B-D) Plots showing the average density of GABAergic synapses, GAD65+ puncta, or gephyrin+ puncta on proximal dendrites of 16 DIV hippocampal neurons, normalized to the average density of the control condition. $n \geq 45$ neurons per condition from at least three experiments; $p \leq 0.05$, Kruskal-Wallis test. Statistical differences are indicated by different letters and bars sharing a letter are not statistically different. Error bars are ± standard error of the mean. Note that the control condition in some cases does align with the dotted line at 1 due to mathematical outlier removal; see methods.
